# Association of gut microbiota with cerebral cortex and cerebrovascular abnormality in human mild traumatic brain injury

**DOI:** 10.1101/2020.07.19.211227

**Authors:** Lijun Bai, Tianhui Li, Ming Zhang, Shan Wang, Shuoqiu Gan, Xiaoyan Jia, Xuefei Yang, Yinxiang Sun, Feng Xiong, Bo Yin, Yi Ren, Guanghui Bai, Zhihan Yan, Xin Mu, Feng Zhu

**Author notes:** This author supervised this work: F. Z. These authors contributed equally to this work (Lijun Bai, Ming Zhang).

## Abstract

Key roles of the gut–brain axis in brain injury development have been suggested in various mouse models; however, little is known about its functional significance in human mild traumatic brain injury (TBI). Here, we decipher this axis by profiling the gut microbiota in 98 acute mild TBI patients and 62 matched controls, and subgroup of them also measured circulating mediators and applied neuroimaging. Mild TBI patients had increased α-diversity and different overall microbial compositions compared with controls. 25-microbial genus classifiers distinguish patients from controls with an area under the receiver operating characteristic curve (AUC) of 0.889, while adding serum mediators and neuroimaging features further improved performance even in a small sample size (AUC = 0.969). Numerous correlations existed between gut bacteria, aberrant cortical thickness and cerebrovascular injury. Co-occurrence network analysis revealed two unique gut–brain axes in patients: 1) altered intestinal *Lachnospiraceae_NK4A136_group* and *Eubacterium_ruminantium_group*-increased serum GDNF-subcallosal hypertrophy and cerebrovascular injury; 2) decreased intestinal *Eubacterium_xylanophilum_group*–upregulated IL-6–thinned anterior insula. Our findings provide a new integrated mechanistic understanding and diagnostic model of mild TBI.

## Introduction

Traumatic brain injury (TBI) is a public health challenge of vast but insufficiently recognized proportions. It is estimated that more than 50 million people worldwide suffer at least one TBI in any given year, and approximately half the world’s population will have one or more TBIs over their lifetime^1^. TBI is the leading cause of mortality in young adults and a major cause of disability across all ages; although there has been a great deal of researches seeking insights into its pathogenesis, the ongoing pathophysiology of TBI has not been fully elucidated^2, 3^. Recent studies have demonstrated key roles of the microbiome–gut–brain axis in mediating various neuroinflammatory, neurodevelopmental and neurodegenerative diseases^4^. Intestinal dysfunction is highlighted as one of the most common but neglected consequences of TBI^2, 3^. Several psychological and physiological disturbances following TBI are known to be sufficient to alter the balance of gut homeostasis maintained by the microbial ecosystem and host immune system, with results including traumatic stress^5^, dysfunctional intestinal contractility and motility^6, 7^, and increased gut permeability^8^. Accordingly, animal studies have revealed profound changes in gut microbiota after various nerve injuries^6, 7^, such as stroke spinal cord injury^8^ and TBI^4, 5^. More importantly, gut dysbiosis in animal models is not merely a byproduct of injury but an essential contributory factor to TBI-related neuropathology and impaired behavioral outcomes^9^. In contrast to the plentiful evidence from animal studies, the gut microbiome of human patients with TBI has been poorly characterized. Considering the high value of gut microbes in disease diagnosis and prognosis prediction, there is an urgent need to systematically examine the composition and functional capacity of gut microbiota in relation to TBI.

The most severe outcome of TBI in terms of public health comes from long-lasting progressive neurobehavioral sequelae^10^, including a heightened risk of several psychiatric and neurodegenerative diseases^11, 12^. Even mild TBI is one of the strongest environmental risk factors facilitating the development of neurodegenerative diseases, an observation that has been validated widely by population-based studies^12^. For example, mild TBI is a well-established risk factor for a variety of neurodegenerative diseases including Parkinson disease (PD) and dementia, with significant effects that persist for decades after injury^12^. Recently, the regulatory role of gut microbiota in pathophysiology has been revealed with the strongest evidence in PD^13, 14^ and a growing appreciation of its role in Alzheimer’s disease (AD)^15^ and stroke^9, 16^. However, the biological mechanisms linking TBI and increased risk of those neurobehavioral sequelae are still unknown. Longitudinal studies are the best approach to analyze the effects of microbiota on the development of these sequelae; however, such long follow-up periods will postpone the solution of this urgent scientific problem. Another effective method to gain insight into the roles of gut dysbiosis in long-term prognosis is to investigate the relationships between post-injury microbiota alterations and the well-known risk factors/phenotypes related to neurobehavioral sequelae.

Neuroimaging is a rapidly growing technology to noninvasively characterize brain structural and functional alterations in vivo under pathological conditions. Mild TBI accounts for 80-90% of all cases of TBI in both civilian and military populations^17^. Unlike moderate to severe brain injury, mild TBI cannot be diagnosed by conventional CT and MRI. Insight into the neuropathophysiology of mild TBI in humans is mainly dependent on cortical structural and functional changes. Our team, together with other groups, has identified several specific neuroimaging signatures of mild TBI and demonstrated the crucial roles of interhemispheric structural and functional connectivity damage^18^ and alterations in the default-mode network^19^ in chronic brain lesion development after injury. Moreover, accelerated aging processes are exhibited in the brains of patients with mild TBI, including accelerated brain atrophy^20^ and microvascular injury (our unpublished data). Brain atrophy/cortical thinning is a common pathology for multiple neurodegenerative and mental disorders^21, 22^. Brain microvascular injury represents ischemia, hemorrhage, blood–brain barrier (BBB) disruption, local inflammation/immune activation, and neuron death. Cerebrovascular injury in mild TBI has been revealed by both postmortem imaging and histological examination, including hypointensity on T_2_*-weighted MRI (white matter (WM) hyperintensity, WMH) that colocalizes with iron-laden macrophages^23^. These aberrant cerebrovascular changes synergistically interact with neurodegenerative pathologies, such as decreased cortical thickness, lowering the threshold for AD^24–27^. Neuroimaging is currently the most important technology to characterize brain function in living human subjects; therefore, we focus on the interaction between gut microbiota and neuroimaging phenotypes, which is key to understanding the functionality of the human microbiota–gut–brain axis. Primary evidence has also shown that the human gut microbiome profile is significantly associated with cerebrovascular dysfunction and brain structure. Regarding the cortical thinning and microvascular injury involved in mild TBI, the modulatory effects of the gut microbiome on these features are still unclear. Thus, we sought to characterize the gut microbiome signature of individuals with mild TBI and identify gut microbes associated with two common neural phenotypes involved in both brain injury and its long-term neurobehavioral sequelae/comorbidities, i.e., cortical morphology^28^ and microvascular injury^23^.

Here, we profiled the gut microbiota of 98 patients with mild TBI and 62 well-matched healthy controls (HCs) via 16S rRNA sequencing. To identify specific alterations in the gut–brain axis, we also compared potential circulating mediators and brain structural and functional traits between patients and controls. The relationships among dysbiotic gut microbiota, 6 increased serum molecules that may link the gut and brain, aberrant cerebral morphology and microvascular injury (WMH), and symptom severity (Rivermead Post-Concussion Symptom Questionnaire, RPCS) were analyzed to identify the gut–brain axis abnormalities underlying mild TBI. Based on these alterations in the gut–brain axis, we further constructed several diagnostic models that performed very well in discriminating patients from controls.

## Results

### The gut microbiota profile of mild TBI patients

A total of 98 patients with mild TBI and 62 matched HCs were included in this study. There were no significant differences in demographic or clinical characteristics between these two groups (*P* > 0.05; Table 1; detailed data in Supplementary Data 1). The gut microbiota of mild TBI patients exhibited increased α-diversity at the genus level, including the Chao1 and abundance-based coverage (ACE) indices, compared with that of HCs (*P* = 0.031 and 0.011, respectively; Wilcoxon rank-sum test shown in Fig. 1a and b). Additionally, the comparison of β-diversity indexes based on both the unweighted UniFrac distance and Bray-Curtis dissimilarity indicated that the overall microbiota composition of patients with mild TBI varied markedly from that of HCs (all *P* = 0.001, permutational multivariate analysis of variance (PERMANOVA); Fig. 1c and Supplementary Fig. 1). At each taxonomic level from phylum to genus, we identified numerous classical taxa that were differentially enriched in mild TBI patients (Supplementary Data 2). At the phylum level, the abundance of Firmicutes, one of the most dominant phyla in the healthy gut, was markedly decreased in the mild TBI group (*P* = 0.011, Wilcoxon rank-sum test; Fig. 1d and e). At the genus level, linear discriminant analysis (LDA) effect size (LEfSe) analysis revealed that 72 bacterial taxa with LDA scores > 2.0 and 15 bacterial taxa with LDA scores > 3.0 displayed statistically significant differences in relative abundance between the patients and the controls after adjusting for age, gender, body mass index (BMI), smoking, drinking and bowel habits (*P* < 0.05; Fig. 1f; Supplementary Data 3). Compared with controls, 15 differentially enriched genera with LDA scores > 3.0 were identified in the gut of patients with mild TBI, including 8 upregulated bacteria, namely, *Escherichia_Shigella*, *Ruminococcus_gnavus_group*, *Bifidobacterium*, *uncultured_f_Muribaculaceae, Lactobacillus*, *Clostridium_innocuum_group*, *Catenibacterium*, *and Alloprevotella*, and 9 downregulated bacteria, namely, *Prevotella_7*, *Dorea, Dialister*, *Eubacterium_ruminantium_group*, *Fusicatenibacter*, *Lachnospiraceae_NK4A136_group*, *Fusobacterium*, *Agathobacter*, and *Faecalibacterium*. Next, co-occurrence correlations between 72 mild TBI-associated gut microbial genera at LDA score > 2.0 were shown in a network (Supplementary Fig. 2). Intriguingly, the genera enriched in mild TBI were more interconnected than those enriched in controls (ρ < −0.4 or > 0.4, *P* < 0.05). A relatively isolated network including 21 genera was present in the gut of patients; this network was characterized by extensive strong positive associations (ρ > 0.6) among the genera within the network and a few weak associations between the genera inside and outside the network. The representative bacteria in this relatively isolated network comprised *Anaeromyxobacter*, *Bradyrhizobium*, *Desulfatiglans*, *Eubacterium*, *Sphingomonas*, *Sulfuricurvum*, *Thiobacillus*, *Ochrobactrum*, and others (Supplementary Fig. 2).

**Fig. 1.**
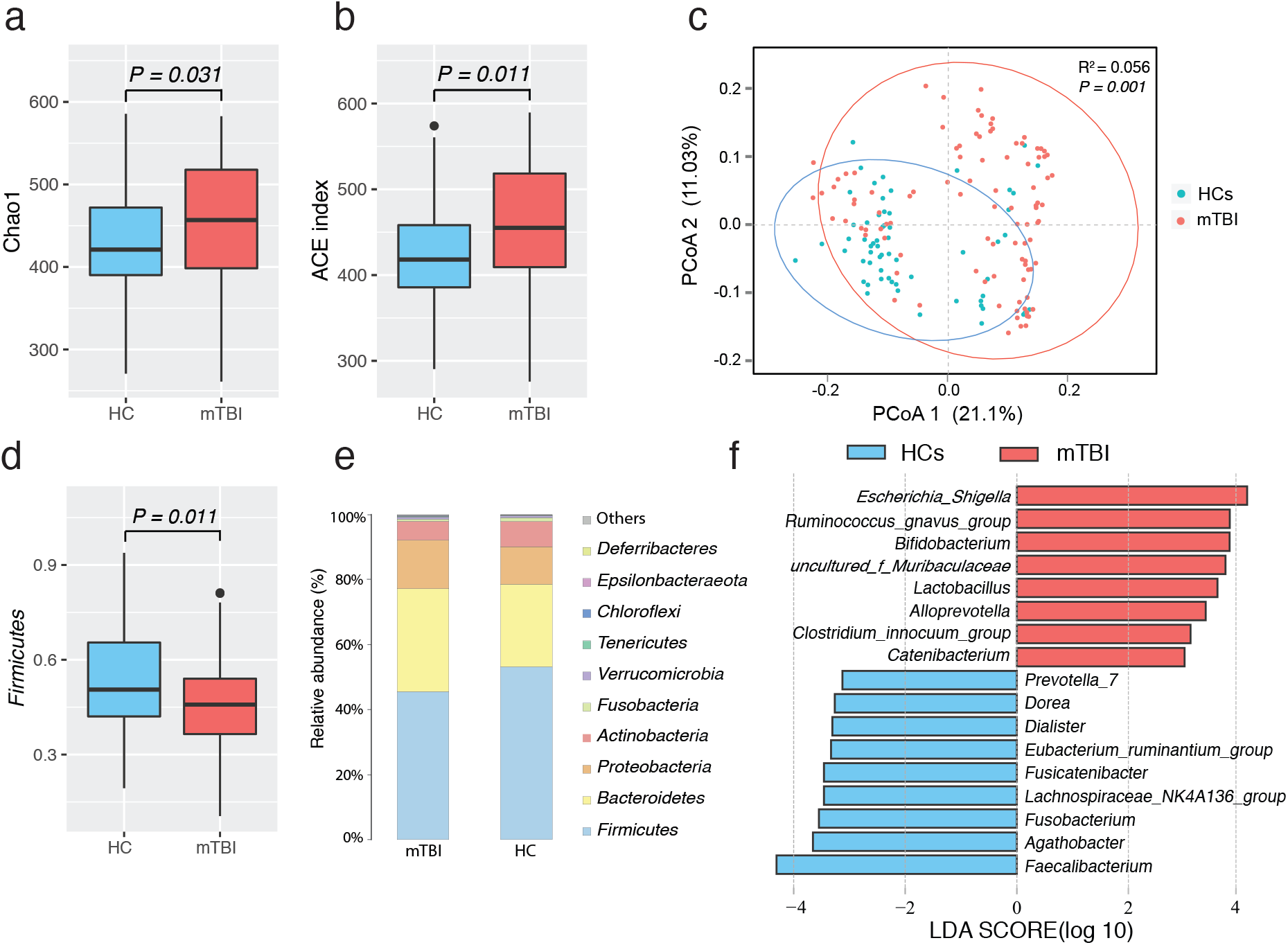
Comparisons of α-diversity and β-diversity between patients with mTBI (n = 98) and HCs (n = 62). The number of Chao1 index **(a)** and ACE index **(b)** were significantly increased in mTBI compared with controls. *P* values were calculated via the Wilcoxon rank-sum test. **(c)** PCoA based on the unweighted UniFrac matrix showed that the overall fecal microbiota composition was different between patients and controls (*P* = 0.001). *P* values were calculated via the PERMANOVA test. **(d)** Firmicutes were significantly decreased in mTBI compared with controls. *P* values were calculated via the Wilcoxon rank-sum test. **(e)** Relative proportions of bacterial phyla are presented in mTBI and controls. **(f)** LDA effect size analysis revealed that the relative abundance of 17 genera was significantly different between mTBI and controls. Boxes represent the medians and interquartile ranges (IQRs) between the 25th and 75th percentiles; whiskers represent the lowest or highest values within 1.5 times IQR from the first or third quartiles. mTBI, mild traumatic brain injury; HCs, healthy controls; PCoA, principal coordinate analysis; LDA, linear discriminant analysis. See detailed statistical data in Supplementary Source Data file.

### Functional potential of the gut microbiota in mild TBI

Functional modules and pathways enriched in the gut microbiota of patients compared with controls were analyzed using the Kyoto Encyclopedia of Genes and Genomes (KEGG) database (Supplementary Data 4). We screened out the KEGG categories that were differentially enriched by the gut microbiota of patients via PICRUSt. Statistical Analysis of Metagenomic Profiles (STAMP) analysis detected 19 KEGG pathways and modules that were significantly different between mild TBI patients and controls (*P* < 0.05, false discovery rate (FDR) < 0.05). Briefly, mild TBI-depleted microbial functional modules included insulin resistance, epithelial cell signaling in *Helicobacter pylori* infection and peptidoglycan biosynthesis, whereas mild TBI-enriched functional modules included type I polyketide structures, bile secretion, the synaptic vesicle cycle, retrograde endocannabinoid signaling, lipoic acid metabolism, and others. (Fig. 2a). Next, we predicted the alterations in microbial high-level phenotypes using BugBase^29^. Compared with the controls, facultatively anaerobic bacteria, gram-negative bacteria and microbial oxidative stress tolerance were upregulated (*P* = 0.01, 0.004, 0.023, respectively) while gram-positive bacteria and anaerobic bacteria (*P* = 0.004, 0.048, resepctively) were downregulated in mild TBI patients (Wilcoxon rank-sum test; Fig. 2b-f).

**Fig. 2.**
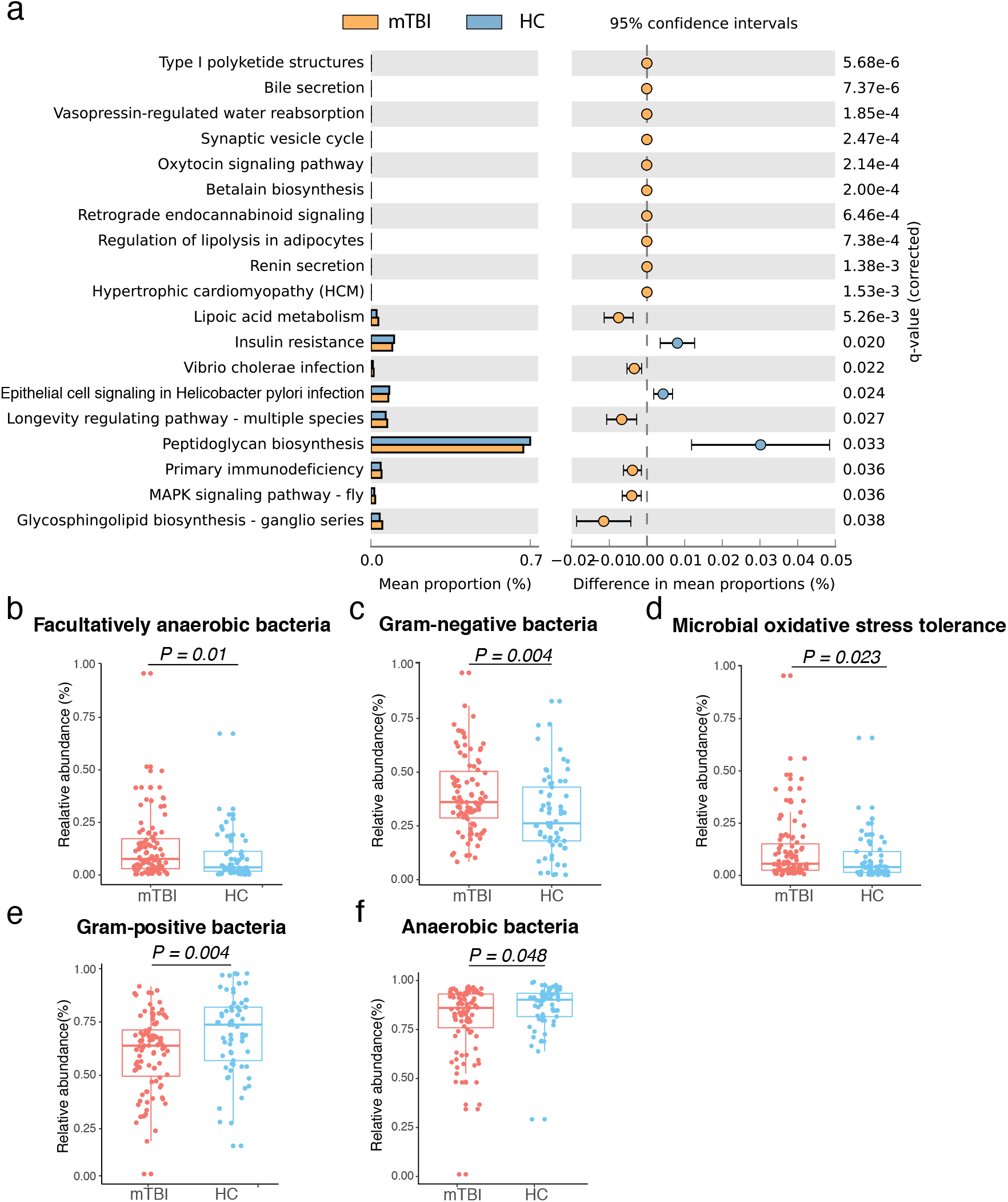
The functional prediction of gut microbiota in mTBI and HCs. **(a)** KEGG Orthology (KO) represented enriched functional pathways between HCs (n = 62) and mTBI (n = 98). **(b-e)** BugBase predicted bacterial community phenotypes in both groups. *P* values were determined by the Mann-Whitney-Wilcoxon test. Boxes represent the medians and interquartile ranges (IQRs) between 25th and 75th percentile; whiskers represent the lowest or highest values within 1.5 times IQR from the first or third quartiles. mTBI, mild traumatic brain injury; HCs, healthy controls. See detailed statistical data in Supplementary Source Data file.

### Gut microbiota signatures of mild TBI

To evaluate the diagnostic potential of the gut microbiome in mild TBI, we constructed a series of random forest disease classifiers based on 72 microbial genera that were differentially enriched between patients and controls. This analysis was conducted with a five-fold cross-validation procedure ten times. A total of 25 genera reached the lowest classifier error in the random forest cross-validation, and the area under the receiver operating characteristic curve (AUC) of this model was 0.895 (95% CI, 0.847-0.944; Fig. 3a). This microbial classifier was not significantly influenced by age, gender, BMI, or diet style (Supplementary Data 5). The traditional biomarkers for mild TBI are a few molecules in blood and cerebrospinal fluid (CSF)^30^. In the present study, we compared between 20 serum biomarkers between mild TBI patients and controls. Of which, 6 serum biomarkers presented significant increases in mild TBI patients after controlling for age, gender and BMI (all for *P* <0.05, Supplementary data 6). To further examine whether the combined gut bacteria and blood serum biomarkers can obtain better diagnostic potential, we screened 72 candidate molecules and 6 increased serum biomarkers with differential abundance in patients compared with controls entered into the random forest model. Finally, a 20-factor classifier including 14 microbial genera and 6 serum markers fulfilled the lowest classifier error in the random forest cross-validation, with an AUC of 0.954 (95% CI, 0.917-0.991; Fig. 3b), which also displayed a better diagnostic performance than the classifier of only 6 serum molecules with AUC of 0.913 (95% CI, 0.859-0.967, Supplementary Fig. 3). The contribution of each gut bacterium or serum molecule to the diagnostic model was shown in Fig. 3c and d. The abundances and concentrations of these gut bacterial genera and serum molecules for the 20-factor classifiers were shown in Fig. 3e and f. The overlaps between the 25-genus classifier and the 14-genus, 6-serum-molecule classifier included 12 bacterial genera: *Eubacterium_ruminantium_group, Ruminiclostridium*, *Ruminococcus_1, uncultured_f_Muribaculaceae, Psychrobacter, Fusobacterium, Ruminococcaceae_UCG_003, Sphingomonas, Bradyrhizobium*, *Catenibacterium*, *uncultured_c_Anaerolineae*, and *Rubus_hybrid_cultivar*.

**Fig. 3.**
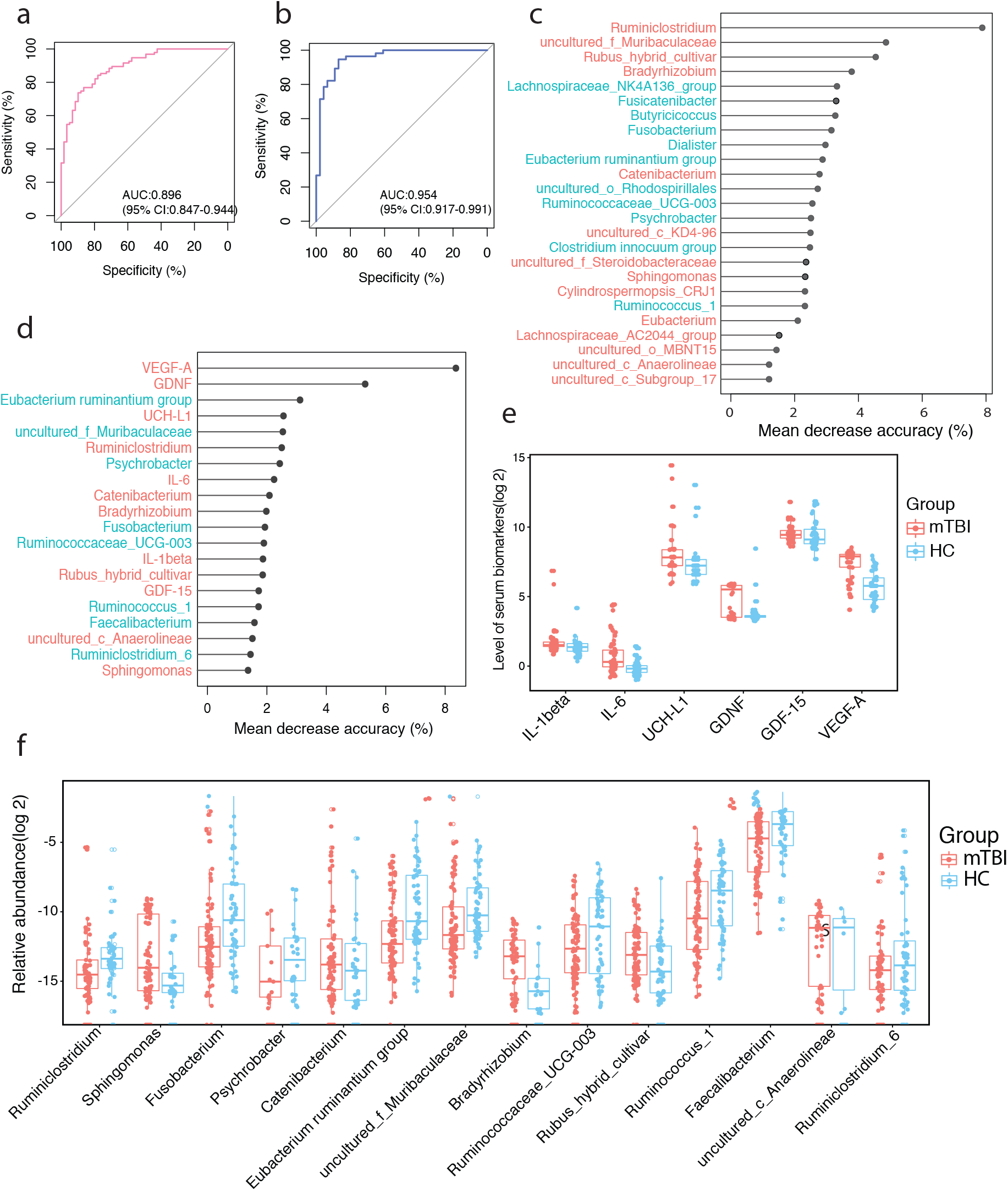
Gut microbiome and serum molecule-based discrimination between mTBI patients and HCs. (**a)** Receiver operating characteristic curve (ROC) based on 98 mTBI patients from 62 HCs was calculated by cross-validated random forest models. The area under the ROC curve (AUC) and the 95% confidence intervals (CI) are also shown. **(b)** The combination of both gut microbiota and blood serum biomarkers as classifiers was selected by cross-validated random forest models to discriminate 56 patients from 46 controls. The AUC and the 95% confidence intervals are also shown. **(c)** The 25 forest models.The length of the line indicates the contribution of the genus to the discriminative model. The color of each genus indicated its enrichment in mTBI patients (red) or HCs (blue). **(d)** The 6 serum biomarkers and 14 genera with the most weight to discriminate mTBI and HCs were selected by cross-validated random forest models. The length of the line indicates the contribution of the genus to the discriminative model. The color of each genus indicated its enrichment in mTBI patients (red) or HCs (blue). The relative level (log2) of 14 genus abundance **(e)** and 6 serum biomarker **(f)** classifiers used in the diagnosis model **(d)**. Each dot represents one value from an individual participant, and the boxes represent the medians and interquartile ranges (IQRs) between the first and third quartiles; whiskers represent the lowest or highest values within 1.5 times IQR from the first or third quartiles. Outliers are not shown. mTBI, mild traumatic brain injury; HCs, healthy controls. See detailed statistical data in Supplementary Source Data file.

### Gut bacteria underlying brain pathology and symptoms

Next, we analyzed the effects of 72 mild TBI-associated bacteria on three types of disease-related phenotypes, i.e., RPCS scores, serum blood biomarkers, cortical thickness and cerebrovascular injury (WMH) (Fig 4a-d). Regarding the RPCS, 16 microbial genera displayed significant associations with symptom severity as measured by total RPCS scores, mainly including *Clostridium_innocuum_group, Hungatella and Sphingomonas* (all for *P* < 0.02, FDR < 0.21, *ρ* > 0.27; Supplementary Data 7; Fig. 4a). Additionally, the abundance of *Clostridium_innocuum_group* and *Sphingomonas* was positively correlated with three subscores of the RPCS, namely, headache, being irritable and fatigue (all for *P* < 0.05, FDR < 0.21, *ρ* > 0.25), while *Lachnospiraceae_NK4A136_group*, *Eubacterium_ruminantium_group and uncultured_f_Muribaculaceae* were negatively correlated with sleep disturbance and being irritable in mild TBI patients (*P* < 0.05, FDR < 0.28, *|ρ|* >0.23). Next, we compared WMH and cortical thickness between mild TBI patients and controls. Total WMH volumes displayed nominally significant increases in mild TBI patients compared with HCs after controlling for age, sex, education and ICV (intracranial volume, brain plus associated CSF with the inner table of the skull as the outer boundary of the segmented image) (*F*_5, 40_ = 3.772, *P* = 0.059; Fig. 4e). Increased WMH volumes were also identified in the frontal and temporal lobes of mild TBI patients (*F*_5, 40_ = 4.74 and 6.31, *P* = 0.036 and 0.016, respectively; Fig. 4f and g). With respect to brain morphological changes, we found that mild TBI patients had markedly decreased cortical thickness in the right anterior insula and left hippocampus/parahippocampus (Hipp/PH) and increased cortical thickness in the right subcallosal compared with controls after adjusting for age, sex, education and ICV (F_1, 54_ = 10.24, 5.22, 4.74, all for *P* < 0.05, FDR corrected; Fig. 4h-j). Next, partial Spearman’s rank-based correlation analysis was conducted to analyze the associations of gut microbiota with 6 serum molecules and 6 neuroimaging features exhibiting significant between-group differences, controlling for age, gender, BMI, education, smoking, drinking and bowel habits (Supplementary Data 8). A total of 26 microbial genera displayed significant correlations with at least one type of disease-related phenotype and were mainly present in the group of patients (Fig. 4h-j). Intriguingly, six genera displayed associations with multiple phenotypes (for both serum molecules and neuroimaging features). *Lachnospiraceae_NK4A136_group* associated with RPCS symptoms and exhibiting diagnostic potential significantly correlated with not only GDNF but also the right subcallosal and frontal WMH (*P* = 0.00008, *ρ* = 0.528; *P* = 0.005, *ρ* = −0.685; *P* = 0.021, *ρ* = 0.608). The *Eubacterium_ruminantium_group* associated with RPCS symptoms and diagnostic potential was negatively correlated with the right subcallosal thickness and positively correlated with GDNF (*P* = 0.003, *ρ* = −0.805; *P* = 0.001, *ρ* = 0.45, respectively). The *Clostridium_innocuum_group* associated with RPCS symptoms and diagnostic potential was also positively associated with the left Hipp/PH thickness and negatively associated with GDNF (*P* = 0.01, *ρ* = 0.641; *P* = 0.009, *ρ* = −0.365, respectively). *Achromobacter* was positively correlated with VEGF-A and subcallosal thickness (*P* = 0.006, *ρ* = 0.386; *P* = 0.022, *ρ* = 0.587, respectively). *Eubacterium_xylanophilum_group* was negatively correlated with IL 6 and cortical thickness of the right anterior insula (*P* = 0.042, *ρ* = −0.289; *P* = −0.041, *ρ* = −0.532). *Robinsoniella* was negatively associated with IL 6 and positively related with cortical thickness in the left Hipp/PH (*P* = 0.044, *ρ* = −0.286; *P* = 0.037, *ρ* = 0.541).

**Fig. 4.**
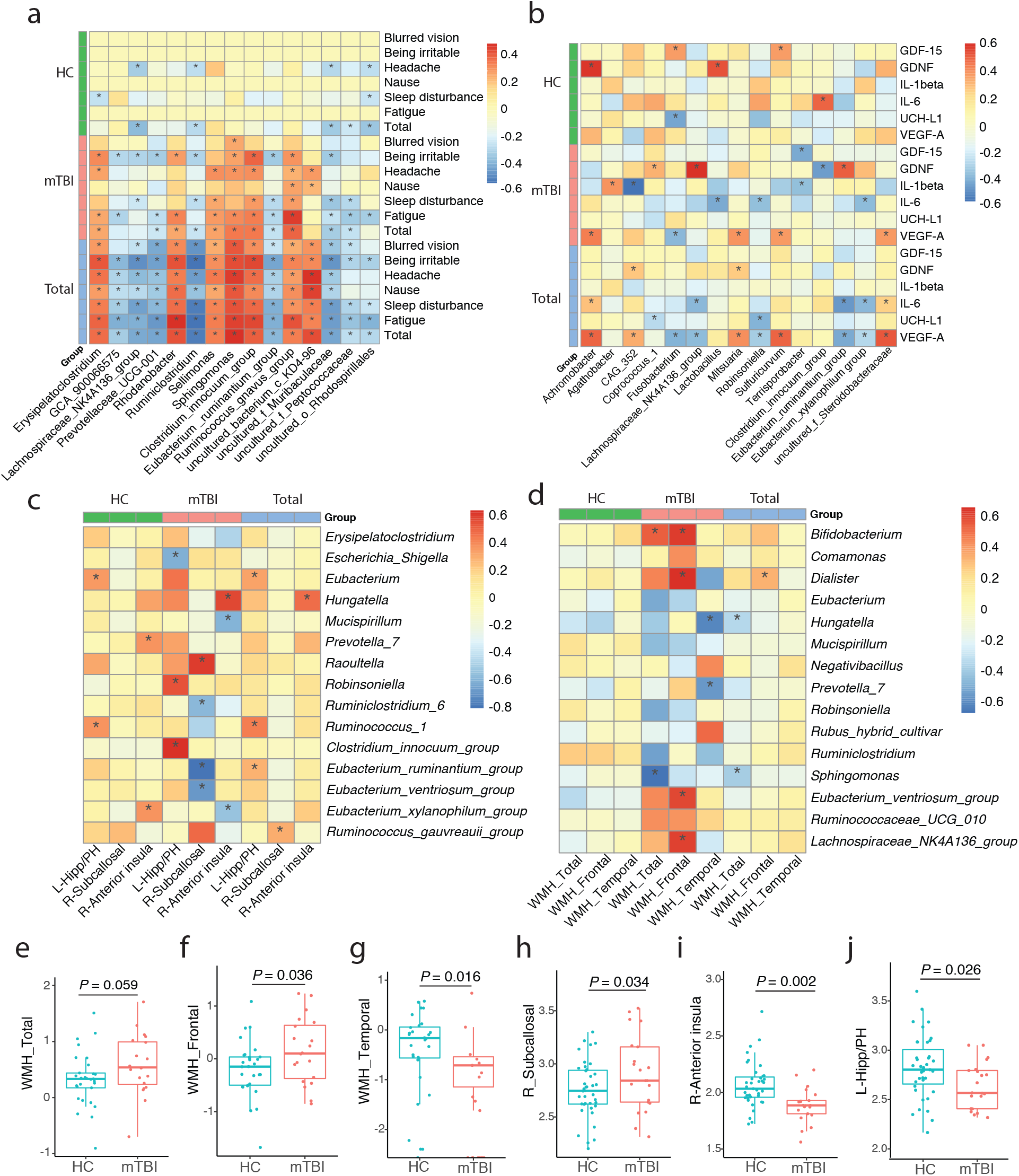
Associations between gut microbiota, RPCS, blood serum molecules and brain features in mTBI. Patients (n = 20) presented a nonsignificant difference in the total WMH **(a)** and a significantly higher WMH in the frontal lobe **(b)** and parietal lobe **(c)** compared with HCs (n = 39). One-way analysis of variance (ANOVA) indicated that patients showed increased cortical thickness in the right subcallosal gyrus **(d)** and decreased cortical thickness in the right anterior insula **(e)** and left hippocampus/parahippocampus (Hipp/PH) **(f)**. Partial Spearman correlation analysis was first conducted between 72 genera (LDA > 2.0) and total (or subitem) RPCS **(g)**, serum biomarkers **(h)**, cortical thickness **(i)** and WMH **(j)** in the mild TBI, healthy controls and total subject groups. Only the top 15 genera with higher relation coefficients for each correlation type were finally presented on the heatmap. * indicates *P* < 0.05. mTBI, mild traumatic brain injury; HCs, healthy controls. See detailed statistical data in Supplementary Source Data file.

### Two associations between gut bacteria and brain aberrations in mild TBI

Next, we explored the relationships among gut microbiota, circulating molecules, and brain structural and functional abnormalities via a co-occurrence correlation network (Fig. 5a). We define a gut–brain axis as a series of co-occurrence correlation networks that link gut bacteria, serum mediators, and cerebral traits (denoted as nodes) (Fig. 5b). We defined the node degree as a metric to quantify the number of edges (i.e., connections) connected to that node. For both serum mediators and cerebral traits, important node hubs were then defined as node degree > 3, including GDNF, VEGF-A, and IL 6 in serum mediators, as well as the anterior insula thickness, subcallosal thickness and frontal WMH in cerebral traits. Finally, two axes were identified: 1) a gut bacteria–GDNF–subcallosal thickness–WMH axis and 2) a gut bacteria– IL-6–anterior insula axis. In the gut bacteria–GDNF–subcallosal thickness–WMH axis, mild TBI-associated *Clostridium_innocuum_group*, *Lachnospiraceae_NK4A136_group*, and *Eubacterium_ruminantium_group* were related to post-concussion symptom severity (all for *P* < 0.05) and serum GDNF level (all for *P* < 0.01). *Lachnospiraceae_NK4A136_group* and *Eubacterium_ruminantium_group* were also associated with cortical thickness of the subcallosal area (all for *P* < 0.005). Moreover, circulating GDNF correlated with not only three gut bacteria but also total WMH and subcallosal cortex thickness (*P =* 0.022, *ρ* = 0.568; *P* = 0.011, *ρ* = −0.599, respectively). In the gut bacteria–IL-6–cortical thickness axis, *Eubacterium_xylanophilum_group* negatively correlated with both IL-6 and cortical thickness of the right anterior insula, and accordingly, IL-6 also correlated with cortical thickness of the right anterior insula (*P* = 0.046, *ρ* = 0.489). Next, we sought to determine whether these two axes are specifically presented in patients with mild TBI. We found that these co-occurrence correlations among gut bacteria, serum mediators, and cerebral traits in each axis were not significant in controls. 6 classifiers with top 40% important weights in the diagnose model, 5 neuroimaging features that varied markedly between patients and controls, and/or 5 gut genera displayed associations with both serum molecules and neuroimaging features were further entered into LASSO regression analyses, in order to explore the diagnose potential of mild TBI from HCs. Finally, a 6-factor classifier including 1 microbial genera, 3 serum markers and 2 neuroimaging features provided a good identification of mild TBI from HCs with an AUC of 0.969 (95% CI, 0.928-1; Supplementary Fig. 4).

**Fig. 5.**
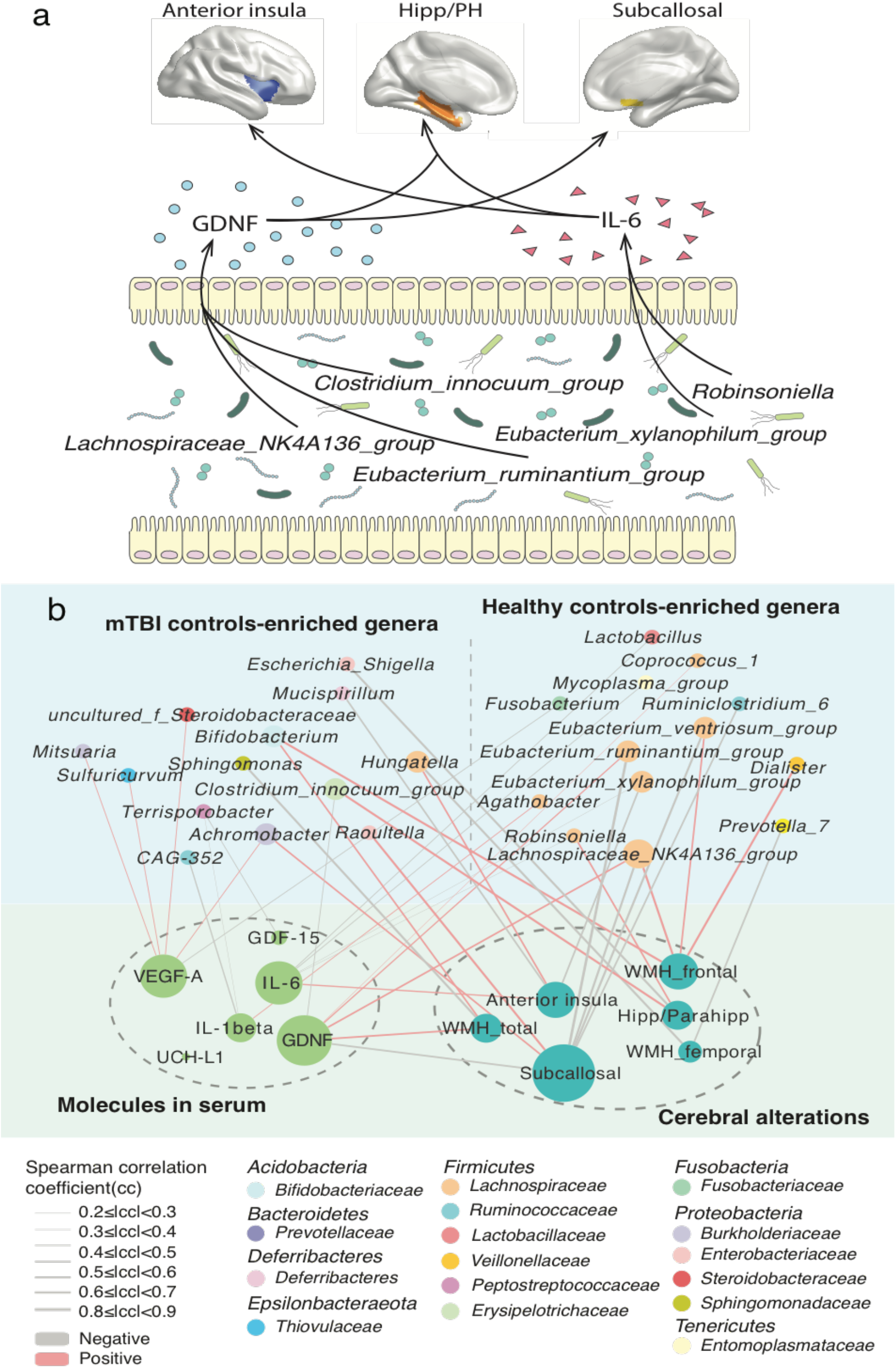
Co-occurrence network of gut microbiome, serum molecules and brain alterations in mild TBI patients. **(a)** An illustration map of their associations. **(b)** Associations of gut genera with serum molecules and/or brain alterations. Node size represented the node degree denoting as the number of edges (i.e., connection) connected to that node. The color of the edge represents positive (red, rho > 0.2, *P* < 0.05) and negative (gray, rho < −0.2, *P* < 0.05). Hipp/PH, hippocampus/parahippocampus; mTBI, mild traumatic brain injury. See detailed statistical data in Supplementary Source Data file.

## Discussion

Here, we integrated three types of datasets from two terminals of the gut–brain axis, i.e., the gut microbiome and neuroimaging traits, and from certain circulating mediators as important regulators to implicate potential mechanistic links in the gut–brain axis of mild TBI. For the first time, we identified two unique altered gut–brain axes in mild TBI associated with serious symptoms. Previous studies on animal models^31^ and human studies^32^ have mainly focused on alterations in the brain or biomarkers and have failed to enrich our understanding of the systems biology underlying mild TBI^33^. Our findings outlined more integrative pathological mechanisms underlying mild TBI. Over the past five years, numerous dysbiotic gut bacteria have been identified in neuropsychiatric conditions, but alterations in the pattern/mode of gut– brain communication are rarely explored in humans. The extensive co-occurrence correlations between dysbiotic bacteria, circulating mediators, and abnormal brain traits suggested that pathway-level changes indeed exist in the gut–brain axis following TBI.

Systemic communications between the gut and brain are regulated by some mediators in the blood, such as immune modulatory metabolites, gut peptides, neurotransmitters, and cytokines^34^. We screened 6 mild TBI-associated serum molecules from 20 candidates, some of which are accepted as serum biomarkers for TBI^35, 36^. Of these 6 molecules, GDNF and IL-6 were identified as key hub nodes linking dysbiotic gut microbiota and aberrant brain traits in a co-occurrence network and marked two unique alterations in the gut–brain axis. The altered axis of gut bacteria–GDNF-subcallosal thickness and WMH included two differentially regulated gut bacteria (*Lachnospiraceae_NK4A136_group* and *Eubacterium_ruminantium_group*), increased serum GDNF, and thicker subcallosal area in patients with mild TBI. The subcallosal area is one of largest neurogenic regions in the adult brain. Our results revealed, for the first time, that the subcallosal gyrus became thickened within 5 post-injury days (PID)^2–5^ following mild TBI, consistent with previous mouse animal evidence that posttraumatic gliogenesis in the adult brain is contributed by progenitor populations in the subcallosal area^37^. Although the functional roles of subcallosal gyrus thickening in TBI development are still unclear, the strongly positive correlation between subcallosal thickness and symptom severity (r = 0.64) suggested that hypertrophy of this area was implicated in brain pathology following mild TBI. Downregulated *Lachnospiraceae_NK4A136_group* and *Eubacterium_ruminantium_group* after injury were positively associated with GDNF levels. The negative associations of GDNF and these two bacteria with subcallosal hypertrophy further implicated that post-injury increased GDNF secretion and bacterium proliferation enhanced the inhibition of cortical thickening in this brain region, which relieved symptom severity.

Moreover, these gut bacteria may modulate the subcallosal gyrus by influencing WM myelination and cerebrovascular injury. The subcallosal area is situated below the genu of the corpus callosum (CC), and WM myelination of the genu of the CC is reported to decline as visible cerebrovascular injury increases in the form of WMH and captures individual variability in systemic vascular health^38^. Disorganized axonal interconnections in efferent pathways from the frontal white matter can also modulate the cortical thickness of the subcallosal area^39^. WMH may provide an additive “second hit” impact on neural transmission along the white matter pathways, which affects the cortical thickness of the subcallosal area. In this way, TBI can be considered a trigger, as well as a useful model to understand certain pathological features of neurological disorders, since it is apparent that the combination of vascular pathology and neurodegenerative changes (i.e., cortical thickness alteration) is additive, lowering the threshold of dementia risk^40^. In the present study, we also found elevated VEGF-A levels in acute mild TBI, and upregulated *Achromobacter* can further predict higher levels of VEGF-A and subcallosal hypertrophy. VEGF has a strong capacity to augment neurogenesis and angiogenesis after TBI, especially in vessels^41, 42^. In support of mild TBI instantiating the link with neurodegenerative disorders, intestinal microbes can regulate the core pathology features of both WMH and cerebral morphological changes underlying both neurological disorders and highlight the possibility of therapeutic advances in disease treatment through modulation of the gut microbiota.

Another obvious altered axis of the gut–brain in mild TBI was characterized by downregulated *Eubacterium_xylanophilum_group*, increased serum IL-6 levels, and cortical thinning of the anterior insula. The anterior insula serves as the primary cortical destination for afferent interoceptive signals from the entire gastrointestinal tract^43^. Cortical thinning of the anterior insula has consistently been reported in many chronic inflammatory and painful visceral conditions, including inflammatory bowel diseases and irritable bowel syndrome^44, 45^. Our previous study indicated that systemic inflammation activation occurs in patients with mild TBI in the acute phase of injury and last for the 3-month follow-up period^32^. Therefore, the thinned anterior insula after mild TBI may be due to a dysfunctional inflammatory response to injury. Here, we further demonstrated that increased serum IL-6 was positively correlated with the cortical thickness of the anterior insula, suggesting that promoted IL-6 secretion is a protective factor to suppress this cortical atrophy. The gut *Eubacterium xylanophilum group* decreased after injury and was negative for IL-6 and anterior insula thickness. It is indicated that the more this bacterium decreased, the higher serum IL-6 and the thicker anterior insula, and the less serious mild TBI symptoms. There are also three other bacteria associated with serum IL-6, which highlights the modulatory roles of gut microbiota in the immune-brain interactions involved in mild TBI. It is a major challenge to determine the function of gut bacteria in pathological processes, especially in the human gut microbiome. Although animal studies have found some evidence for a disease-causing role of the gut microbiome in brain injury, their applicability to humans is not conclusive^46^. The extensive differences between humans and rodents remind us to be very cautious regarding translating laboratory findings into clinical applications^47^. These two gut–brain axes in mild TBI are preliminarily elucidated by our human data, and future studies need to determine the causal effects between these factors in the gut–brain axis and the underlying biological mechanisms.

To the best of our knowledge, this is the first study to profile gut microbiota in human TBI. Our data indicated that gut dysbiosis was marked at 3-5 days PID. Mild TBI significantly altered the abundances of 7 in 14 phyla, 10 in 22 classes, 26 in 44 orders, 35 in 76 families, and 61 in 205 genera in the gut with FDR < 0.05. By then, a total of 4 published animal studies^48–51^ investigated gut bacteria after TBI and reported that altered gut microbiota emerged at PID 1 and remained significant at PID 3, and gut dysbiosis recovered at PID 7. Accordingly, we selected PID 2-5 as the time of our fecal sample analysis in patients. Rodent studies also found obviously decreased α-diversity of gut microbiota in animals exposed to TBI^49, 50^, which is in contrast to our findings indicating greater α-diversity in patients. Moreover, more differentially enriched phyla, families and genera were found in the present study of human TBI than those identified by the rodent model of TBI. These discrepancies in findings can be attributed to the intrinsic differences in gut microbiota and host physiology between humans and animals, as well as to the varied exposure factor (mild vs. moderate-severe TBI) or sampling location (feces vs. cecum/jejunum). Some consistent findings also existed between our human study and previous animal studies, for example, decreased *Firmicutes* and *Deferribacteres* and increased *Bacteroidetes* and *Proteobacteria*^49, 50^ in phylum, increased *Enterobacteriaceae* family^49^, decreased *Eubacterium ventriosum* and increased *Clostridiales* and *Eubacterium* genera^51^. These overlapping bacterial alterations between injured humans and animals suggest that similar bacteria-modulating effects are exerted by common pathophysiological processes underlying human and rodent TBI.

One important application of the microbial signature of a given disease is to develop objective diagnostic markers. Identifying microbial diagnostic markers is especially crucial for mild TBI, as the traditional diagnostic approach is not effective and is dependent on the recall of patients about subjective symptoms or CT scans. Notably, all of our patients with mild TBI had negative CT and were very challenging to diagnose clinically. Here, we constructed a diagnostic model of 25 gut bacteria that discriminates patients from controls with an AUC of 0.86. The diagnostic effect of this model is comparable to some well-defined traditional serum biomarkers of mild TBI, such as tau, NFL, and GFAP (AUC: ~0.85)^30^. Moreover, we found that a combination of 14 gut bacteria and 6 serum biomarkers further increased diagnostic performance (AUC: 0.954). Therefore, gut bacteria should be considered when adding serum biomarkers to improve diagnostic accuracy. Apart from the diagnostic application, these mild TBI-associated gut bacteria can also be used for translational research. Mild TBI is now a paucity of effective treatments. Two types of probiotics, *Lactobacillus acidophilus* and *Clostridium butyricum*, have been shown to exert neuroprotective effects in mice with TBI^2^. Considering that gut dysbiosis has been shown to contribute to brain injury-related neuropathology and impaired behavioral outcomes in experimental models of animals^49, 51^, we proposed that some TBI-associated bacteria may participate in and even drive some pathophysiological processes. Therefore, differentially presented bacteria in the gut of patients can be candidates for screening future disease-causing bacteria and functional mechanism analysis in animals, as well as therapeutic significance evaluation for future clinical trials. In conclusion, our findings provide a new integrated mechanistic understanding and diagnostic model for mild TBI.

## Methods

### Subject recruitment and clinical assessment

This study was conducted in accordance with the Declaration of Helsinki and approved by the Medical Ethics Committee of The First Affiliated Hospital of Xi’an Jiaotong University (TFAHXJU). It is a publicly registered clinical trial (Identifier: NCT02868671; https://clinicaltrials.gov). All participants signed written informed consent in person. Patients with mild TBI were recruited from patients who had non-contrast head CT because of acute head trauma in the emergency departments of three hospitals, TFAHXJU, the Second Affiliated Hospital and Yuying Children’s Hospital of Wenzhou Medical University and Hanzhong Central Hospital. All of mild TBI patients had negative CT scans. The diagnosis of mild TBI was based on the World Health Organization’s Collaborating Centre for Neurotrauma Task Force^52^ as described in our previous study^19^. The inclusion criteria included *i)* Glasgow Coma Score of 13-15; *ii)* one or more of the following: loss of consciousness (if present) < 30 min, posttraumatic amnesia (if present) < 24 h, and/or other transient neurological abnormalities such as focal signs and seizure. Mild TBI participants were excluded following the criteria: *i)* history of neurological disease, long-standing psychiatric condition, head injury, spinal cord injury or a history of substance or alcohol abuse; *ii)* intubation and/or presence of a skull fracture and administration of sedatives on arrival in the emergency department; *iii)* manifestation of mild TBI due to medications by other injuries (e.g., systemic injuries, facial injuries, or intubation) or other problems (e.g., psychological trauma, language barrier, or coexisting medical conditions) or caused by penetrating craniocerebral injury; *iv)* routine laboratory test (blood, urine, stool routine, liver function, renal function) abnormalities, active gastrointestinal diseases or major organic diseases; *v)* the cumulative intake of alcohol greater than 100 ml in the past week; *vi)* antibiotics, probiotics, or prebiotic uses in the past month. HCs did not have any neurological or psychiatric disorders and were well matched to the patients on demographic features (see Supplementary Data 1). Clinical postconcussive symptoms were assessed via the RPCS^53^, which measured the presence and severity of 17 somatic symptoms commonly experienced following head injury (see Supplementary Data 9).

### Fecal Sample Collection and DNA Extraction

Fecal samples from all subjects were collected within 7 acute days postinjury (2.54 ± 1.13 postinjury days) and stored at −80°C within one hour after collection in the hospital. DNA was extracted using the Qiagen QIAamp DNA Stool Mini Kit (Qiagen, Germany) following the manufacturer’s instructions. DNase-free RNase was used during extraction to eliminate RNA contamination. Isolated DNA was quantified using a Qubit 3.0 Fluorometer with a PicoGreen Assay Kit (Thermo Fisher, Shanghai).

### 16S rRNA amplicon sequencing

The V3-V4 region of the 16S rRNA gene was amplified from total fecal DNA using a pair of universal primers as 338F: 5′-ACTCCTACGGGAGGCAGCA-3′ and 806R: 5′-GGACTACHVGGGTWTCTAAT-3′. The PCR volume of 10 μL included 0.2 μL KOD FX Neo, 5 μL KOD FX Neo Buffer, 0.5 μL DNA template (50 ng/mL), 0.3 μL forward prime (10 mmol/L), 0.3 μL reverse prime (10 mmol/L), dNTP 2.0 μL, and the rest volume of ddH_2_O. After an initial denaturation at 95 °C for 5 min, amplification was performed by 25 cycles of incubation for 30 s at 95 °C, 30 s at 50 °C, and 40 s at 72 °C, followed by a final extension at 72 °C for 7 min. Then, the amplified products were purified and recovered using the 1.0% agarose gel electrophoresis method. Library preparation and sequencing were performed on an Illumina HiSeq 2500 Platform with paired-end 250 bp sequences at the Beijing Biomarker Technologies Co., Ltd. (Beijing, China).

### Bioinformatic analysis

All analyses were completed on the Biomarker BioCloud platform (www.biocloud.org) as described in ref.^54^. First, we used FLASH (version 1.2.11; http://ccb.jhu.edu/software/FLASH/) to merge raw reads and then filter high-quality reads by Trimmomatic (version 0.33; http://www.usadellab.org/cms/?page=trimmomatic) and UCHIME (version 8.1; https://omictools.com/uchime-tool). Subsequently, we clustered the denoised tags into operational taxonomic units (OTUs) with similarity ≥97% using USEARCH^55^ (version 10.0; http://www.drive5.com/usearch/) and obtained the OTU unique representative sequences. Taxonomy was assigned to all OTUs by searching against the Silva databases (Release128; https://www.arb-silva.de/) using the RDP classifier within QIIME (version 2.2; http://qiime.org/).

### α-Diversity, β-diversity and functional analysis

α-diversity (Chao1 Index, ACE Index, Shannon’s Index, Simpson index and observed OTUs) was calculated by Mothur^56^ (version 1.30; http://mothur.org/). β-diversity was calculated based on the Bray-Curtis dissimilarity and unweighted and weighted UniFrac metrics using QIIME (version 2.2). Between-group comparisons for α-diversity and β-diversity were conducted using the Wilcoxon rank-sum test and PERMANOVA, respectively. Principal coordinate analysis (PCoA) with unweighted UniFrac distance matrices was performed to ordinate the dissimilarity matrices, and statistical significance was obtained using the PERMANOVA test. The between-group differences in relative abundance were determined by the LDA effect size (LEfSe) pipeline^57^. The 16S RNA sequences were used to impute the metagenomes of the gut microbiome with PICRUSt (Phylogenetic Investigation of Communities by Reconstruction of Unobserved States) as described previously, and the difference between mild TBI and the control was identified with Welch’s t-test in STAMP (v2.1.3)^58^.

### Serum biomarker detection

All blood samples were collected within the same day as the fecal sample. The current serum sample (56 mild TBI patients and 46 matched HCs) included a proportion of participants involved in our previous studies that measured a 9-plex panel of inflammatory cytokines^32^. Details for the collection and analysis of these cytokines can be found in Sun et al^55^. These cytokines included *i)* the archetypal proinflammatory cytokines IL-1β, IL-6, and IL-12 and the anti-inflammatory cytokines IL-4 and IL-10;*ii)* chemokine (C-C motif) ligand 2 or monocyte chemoattractant protein-1 (CCL2 or MCP-1) and member of the CXC chemokine family (CXCL8) IL-8; *iii)* interferon-γ (IFN-γ); and *iv)* tumor necrosis factor α (TNF-α). In the present study, we also included more serum biomarkers, such as *i)* neurotrophins, including brain-derived neurotrophic factor (BDNF), cell line-derived neurotrophic factor (GDNF), GDNF-15, vascular endothelial growth factor A (VEGF-A), β-nerve growth factor (β-NGF); *ii)* neuron-specific enolase (NSE), ubiquitin carboxyl-terminal hydrolase isozyme L1 (UCH-L1), intercellular adhesion molecule (ICAM); *iii)* Park7/DJ, synuclein-α, and IFN-γ.

Serum samples were collected in the morning at 07:00-08:00 h and centrifuged, and aliquots of supernatant were stored at −80 °C until analysis. Serum biomarkers (pg/ml) were measured using reagents on a Luminex multiplex bead system (Luminex Austin, TX, USA). A fluorescence detection laser optic system was used to simultaneously detect binding of each individual protein onto microspheres, thereby allowing analysis of several analyses in a single sample. Intra- and interassay coefficients of variation for Luminex quantification were < 20% and 25%, respectively. Samples with levels that were undetectable by the assay were set to 0.01 pg/ml.

### MRI Data Acquisition

All MRI scans were conducted within 24 h of fecal sample collection. Twenty-one mild TBI patients and thirty-nine demographically matched HCs (named the neuroimaging subgroup) were randomly selected from the whole cohort and received MRI scanning. The MRI (3T GE 750) protocol mainly included the high-resolution T1-weighted 3D MPRAGE sequence (TE = 3.17 ms, TR = 8.15 ms, flip angle = 9°, slice thickness = 1 mm, field of view (FOV) = 256 × 256 mm, matrix size = 256 × 256), and T2 fluid-attenuated inversion recovery (FLAIR; TR = 8000 ms, TE = 94 ms, flip angle = 150°, thickness = 5 mm, FOV = 192 mm × 220 mm, matrix size = 179 × 256). The presence of nonhemorrhagic and microhemorrhagic lesions was independently determined by experienced clinical neuroradiologists (with 9 and 10 years of experience) who assessed multiple modalities of neuroimaging data acquired at baseline (T1-FLAIR; T2-FLAIR; susceptibility weighted imaging, SWI). All subjects were free of any nonhemorrhagic or microhemorrhagic lesions.

### Cortical thickness measurements

The T1-weighted images were then preprocessed to create a 3D model of the cortical surface for further measurements by using the FreeSurfer version 5.2.0 pipeline (http://www.freesurfer.net)^59^. The pipeline included motion correction, nonbrain tissue removal, Talairach transformation, intensity normalization, and white/gray matter segmentation with automatic topology correction^59^. This was followed by registering each subject to a spherical atlas based on parcellation of the cerebral cortex from regions specified by the Destrieux atlas^60^. Regions of interest (ROIs) were selected to include the limbic system (such as the bilateral insula, subcallosal area and Hipp/PH), which are thought to be important in gastrointestinal disorders^61^. The insula was also divided into several subsections according to Destrieux et al^60^. The thickness of each ROI was calculated as the closest distance between the gray-white matter boundary and the pial mesh at each vertex on the tessellated surface^59, 62^. The regional cortical thickness was then calculated as the mean thickness of vertices belonging to ROIs in both hemispheres separately and adjusted for total ICV.

### White matter hyperintensity (WMH) quantification

Cerebrovascular injury is characterized by typical radiological changes on MRI as WMH. WMH quantification was conducted using the T1-weighted and FLAIR images following the Lesion Segmentation Toolbox (LST) pipeline previously described^63^. After data quality control, 20 mild TBI patients and demographically 29 matched HCs from the original neuroimaging subgroup were used for WMH measurements. WMH quantification consisted of the following automated steps: T1-weighted and FLAIR images were skull-stripped and intensity-corrected using the VBM8 toolbox in the Statistical Parametric Mapping (SPM) package. Corrected T1-weighted and FLAIR images were linearly (12-parameter affine) and nonlinearly coregistered. A lesion belief map based on the FLAIR and T1-weighted image was then produced by computing an initial tissue segmentation of the T1-weighted image^63^. This map was refined iteratively weighting the likelihood of belonging to WM or gray matter (GM) against the likelihood of belonging to lesions until no further voxels were assigned to lesions. After thresholding this map with a prechosen initial threshold (k), a lesion map is produced that is subsequently grown along voxels that exhibited hyperintensity in the FLAIR image. The present study set the initial threshold k = 0.3, which is proven to be useful in previous studies^64^. Estimated lesion masks were then automatically filled using an internal filling method proposed by Chard et al^65^. Candidate region voxels were replaced by random intensities from a Gaussian distribution generated from the normal-appearing WM intensities and then filtered to reintroduce the original spatial variation in WM. All imaging analyses were completed without knowledge of demographic and clinical data. The results were also visually inspected for misclassification by one trained rater blinded to the clinical data. A “lobar” atlas was also coregistered linearly to each labeled FLAIR image to define WMH volumes in the frontal, temporal, parietal, and occipital lobes separately^66^. WMH volume was defined as the sum of the labeled voxels multiplied by voxel dimensions; regional volumes were calculated within each labeled lobar region of interest. In an independent cohort of 20 participants, test-retest reliability was greater than 0.98 for both total and regional WMH volumes. Because the distribution of total WMH volume across the population was skewed, it was log-transformed to normalize the distribution. To control for variations in head size^67^, ICV (intracranial volume, brain plus associated CSF with the inner table of the skull as the outer boundary of the segmented image) were also defined using the Brain Extraction Tool (BET) from FSL^68^ with manual modifications performed by trained raters. WMH volumes were calculated in ml, corrected for ICV^69^. ICV were also used as a covariate for further between-group comparison and correlation analysis.

### Construction of the prediction model based on gut microbiome, serum blood molesules and fMRI data

Five-fold cross-validation was performed ten times on a random forest model using the genus abundance profiles of mild TBI patients and HCs. The test error curves from ten trials of five-fold cross-validation were averaged. The classifier model was then chosen based on the minimized sum of the test error and its standard deviation in the averaged curve^70^. The probability of mild TBI was calculated using this set of genera, and a receiver operating characteristic (ROC) curve was drawn (R 3.3.2, pROC package). The correlation between gut bacteria abundance and RPCS scores was calculated by partial Spearman’s rank correlation. Finally, we assessed the possible confounding effects of age, BMI, sex and diet on our random forest model following the procedures of Zeller et al.^70^ (χ^2^ test, Supplementary Data 5).

To futher illustrate the relationship between the gut microbiota, serum blood biomarkers and fMRI data and consider that the disproportion between the samples and parameters used in the prediction model (43 samples and 16 parameters), least absolute shrinkage and selection operator (LASSO) regression was performed to avoid model overfitting^71^. Some variables were eliminated according to penalty rules and potential predictors with non-zero coefficients were removed in LASSO regression^72^. We carried out cross-validation to determined the penalty parameter lambda using the glmnet package (R 3.3.2). We chose the optimal lambda value which minimized the cross-validation error mean and determine the potential parameters^73^. The power of the prediction model was determined by drawing a receiver operating characteristic curve.

### Statistical Analysis

Statistical analyses were performed in SPSS 20.0, and graphs were generated in R (version: 3.6.2). Data are shown as the mean ± standard deviation (SD), mediation and interquartile range, or frequency and percent as indicated. The normal distribution of continuous variables was measured by the Shapiro–Wilk test. The independent two-sample t-test and the Mann-Whitney test were used to compare group differences based on data normality. χ^2^ tests were applied to assess categorical variables. The relative abundance of each genus was compared between the patients and controls via the Wilcoxon rank-sum test followed by Storey’s FDR correction. Only somatic symptoms with prevalence in over 30% of all patients were selected for further correlation analysis with gut microbes. Correlation analyses between the microbiota, RPCS, cortical thickness and WMH data were performed using partial Spearman’s rank-based correlation, controlling for age, sex, education, BMI, smoking, drinking and bowel habits, while the same analysis was applied to the relationship between microbiota and serum molecules by adjusting for age, gender, BMI and dietary habits (R 3.3.2, ppcor package). When analyzing the association between serum blood biomarkers and fMRI data, age, gender, education and BMI were taken into account. In addition, we also conducted surface-based between-group comparisons on specific regional cortical thicknesses using general linear models adjusted for age, sex, education, BMI, smoking, drinking bowel habits and whole-brain mean cortical thickness. *P* < 0.05 was considered statistically significant, and FDR correction was performed for multiple comparisons in all the above analyses.

## Acknowledgements

This study is supported by the National Natural Science Foundation of China (Grant No. 81771914 and 81671671), and Natural Science Foundation of Zhejiang Province (Grant No. LY19H180003).

## Data availability statement

16sRNA sequencing data for donors samples have been deposited in the CNGB Nucleotide Sequence Archive (CNSA) database under accession identification CNP0000119 and in the European Nucleotide Archive (ENA) database under accession identification code ERP111403. The source data underlying all figures except for those not including statistics are provided as a Source Data file.

## Code availability

The following softwares were used: FLASH version 1.2.11, Trimmomatic version 0.33, UCHIME version 8.1, RDP Classifier version 2.2, Mothur version v.1.30, Cytoscape v3.4.0, STAMP v2.1.3. The following R packages were used: ppcor 1.0, ade4 1.7–13, pROC 1.12.1, randomForest 4.6–14.

## Author contributions

L.B. and F.Z: design and conceptualization of the study, interpretation of the data, drafting the manuscript. T.L.: analysis of the data, drafting the manuscript; S. W., S. G, X. J, X. Y, Y. S., F. X. and X. M: analysis and interpretation of the data; B. Y., Y. R., G. B., Z. Y: collecting the data and revising the manuscript for intellectual content; L.B., M.Z. and Z.Y: obtained funding.

## Competing interests

The authors declare no competing financial interests.

## Materials & Correspondence

F.Z. addressed the material requests.

---

**sFig. 1.**
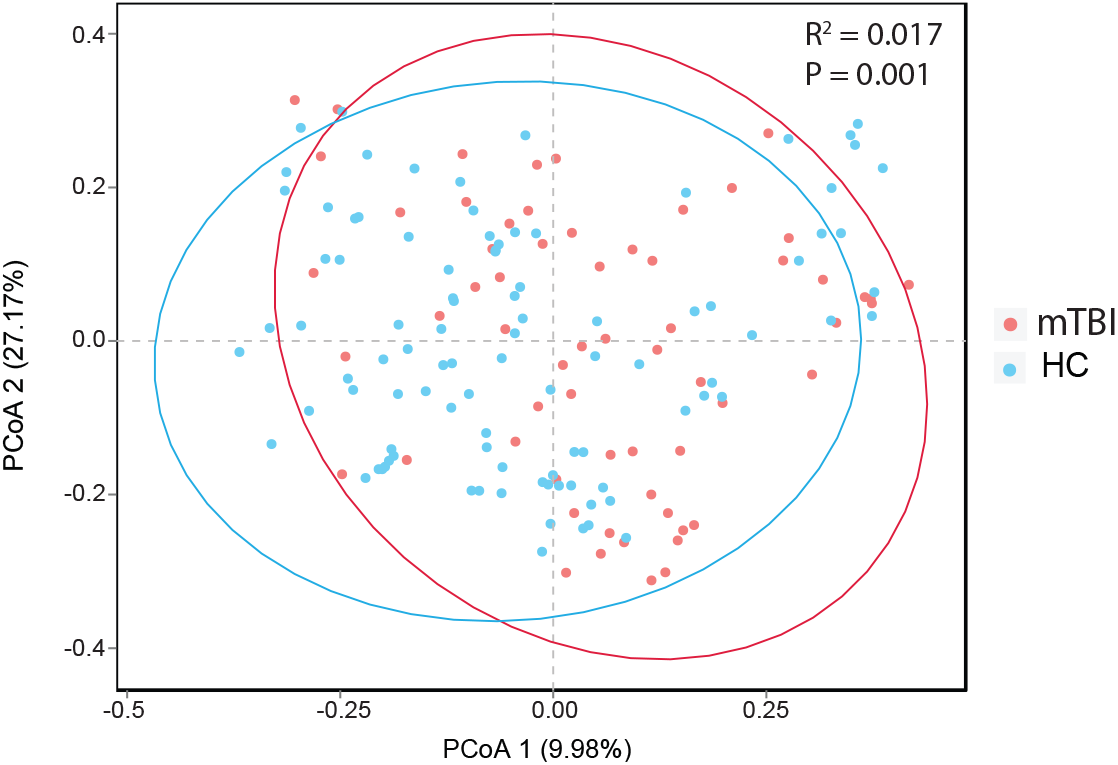
PCoA based on the Bray-Curtis matrix showed that the overall fecal microbiota composition was significantly different between patients and controls (*P* = 0.001). *P* values were calculated by the PERMANOVA test. PCoA, principal coordinate analysis; mTBI, mild traumatic brain injury; HCs, healthy controls. See detailed statistical data in supplementary Source Data file.

**sFig. 2.**
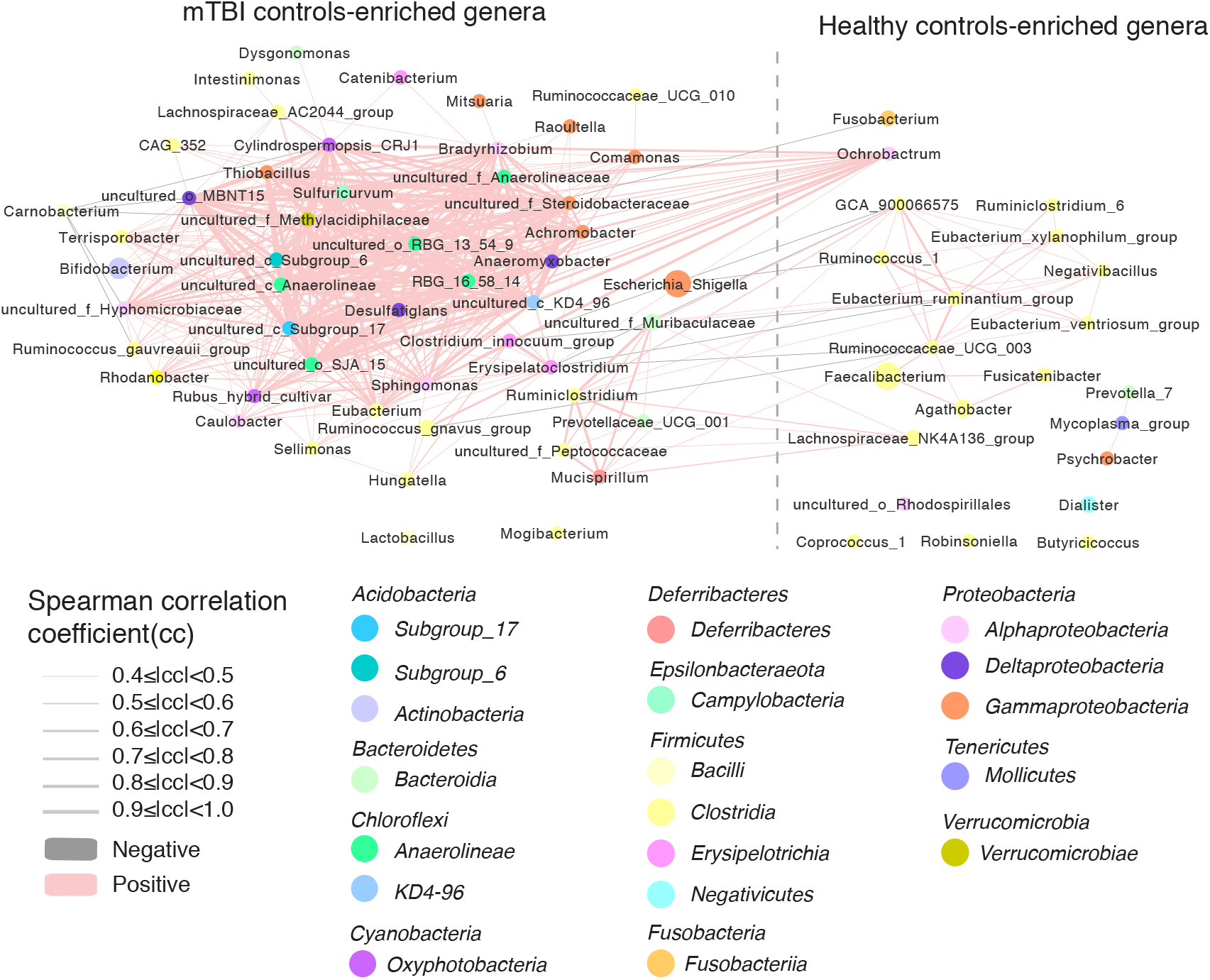
Network of genera differentially enriched in healthy controls and mild TBI patients. Node sizes reflect the mean abundance of significant genera. genus annotated to species are colored according to class (red edges, Spearman’s rank correlation coefficient > 0.4, P < 0.05; blue edges, Spearman’s rank correlation coefficient <−0.4, P < 0.05;). See detailed statistical data in the supplementary Source Data file.

**sFig. 3.**
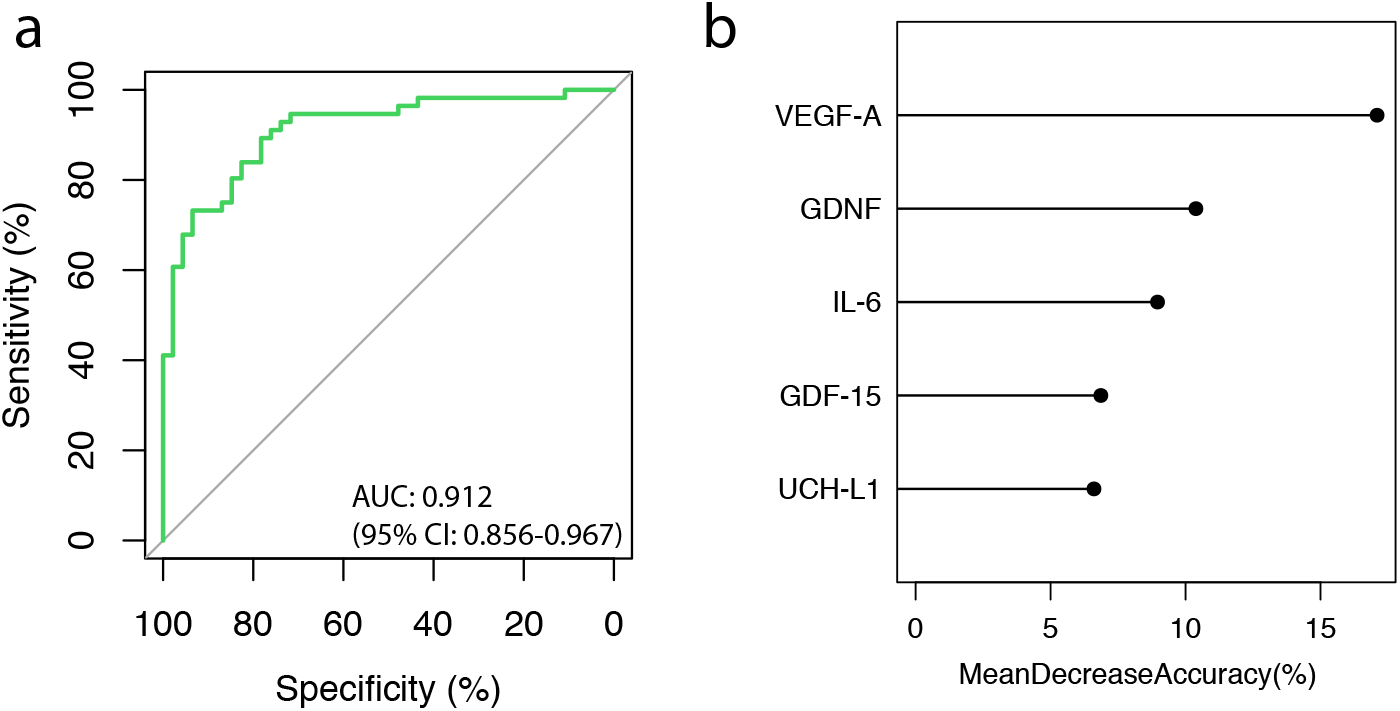
Serum molecule-based discrimination between patients with mild traumatic brain injury (mTBI) and healthy controls. **a.** A classifier containing blood serum biomarkers was selected by the cross-validated random forest models according to 56 patients and 46 controls. The area under the receiver operating characteristic curve (AUC) and the 95% confidence intervals are also shown. **b.** The length of the line indicates the contribution of the serum biomarkers to the discriminative model. See detailed statistical data in supplementary Source Data file.

**sFig. 4.**
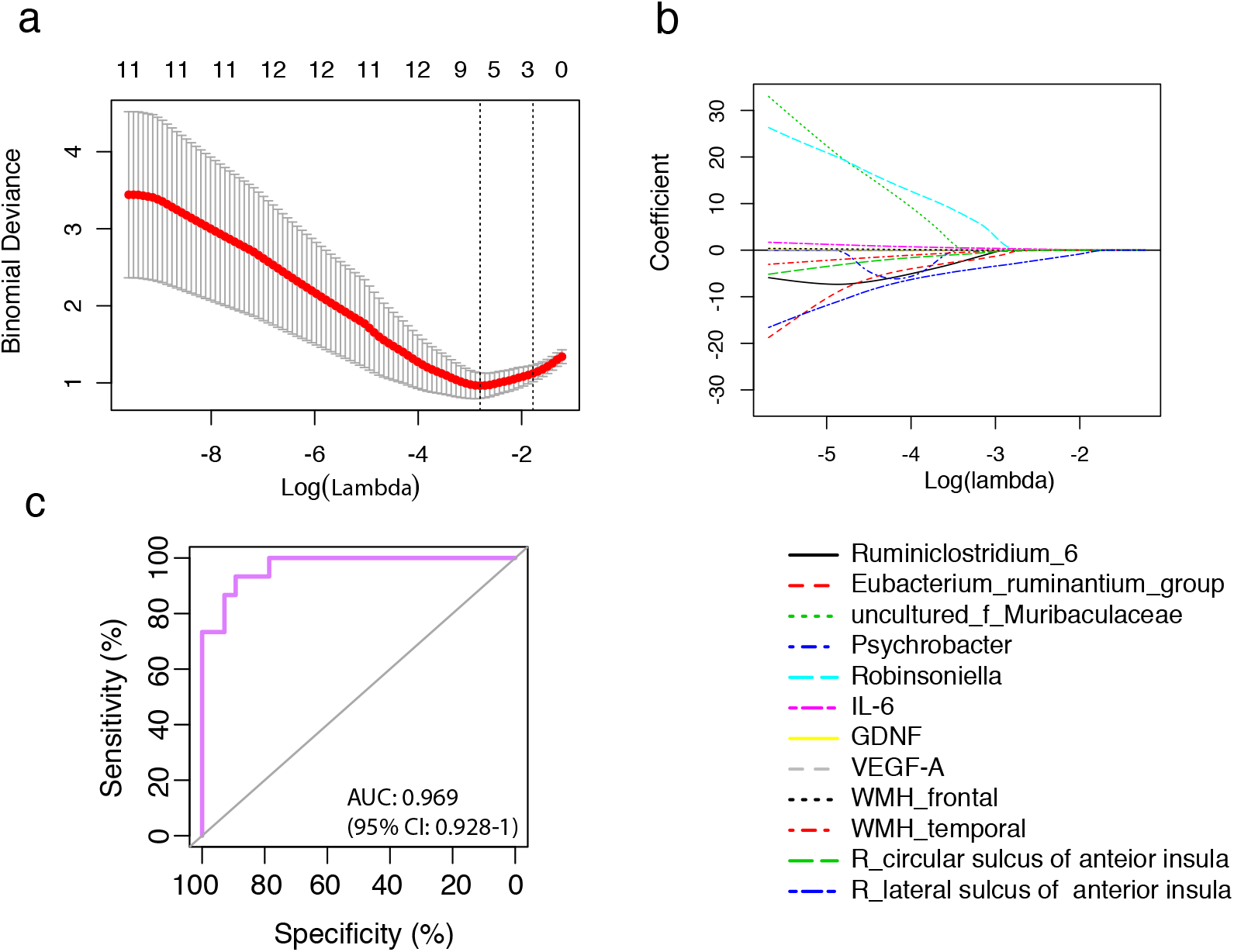
Selection of the optimal parameters used for construction of the optimal prediction model by LASSO regression. (a) Selection of optimal parameter (lambda) in the LASSO model, dotted vertical lines were drawn at the optimal values. (b) LASSO coefficient profiles of the 12 parameters, including 5 generus, 3 serum blood biomarkers and 4 parameters of fMRI data with nonzero coefficients determined by the optimal lambda. (c) The power of the prediction model constructed based no minimum cross-validation error mean was evaluated by Receiver operating characteristic curve (ROC). The area under the receiver operating characteristic curve (AUC) and the 95% confidence intervals are also shown. 15 patients with mild traumatic brain injury and 28 health control were included in this model. See detailed statistical data in supplementary Source Data file.

## Notes

### Competing Interest Statement

The authors have declared no competing interest.

